# Cardiac progenitors auto-regulate second heart field cell fate via Wnt secretion

**DOI:** 10.1101/2021.01.31.428968

**Authors:** Matthew Miyamoto, Suraj Kannan, Hideki Uosaki, Tejasvi Kakani, Sean Murphy, Peter Andersen, Chulan Kwon

## Abstract

Proper heart formation requires coordinated development of two anatomically distinct groups of cells - the first and second heart fields (FHF and SHF). Given that congenital heart defects are often restricted to derivatives of the FHF or SHF, it is crucial to understand the mechanisms controlling their development. Wnt signaling has previously been implicated in SHF proliferation; however, the source of Wnts remains unknown. Through comparative gene analysis, we found upregulation of Wnts and Wnt receptor/target genes in the FHF and SHF, respectively, raising the possibility that early cardiac progenitors may secrete Wnts to influence SHF cell fate. To probe this further, we deleted *Wntless (Wls)*, a gene required for Wnt ligand secretion, in various populations of precardiac cells. Deletion of Wls in Mesp1^+^ cells resulted in formation of a single chamber heart with left ventricle identity, implying compromised SHF development. This phenotype was recapitulated by deleting Wls in cells expressing Islet1, a pan-cardiac marker. Similarly, Wls deletion in cells expressing Nkx2.5, a later-expressed pan-cardiac marker, resulted in hypoplastic right ventricle, a structure derived from the SHF. However, no developmental defects were observed when deleting Wls in SHF progenitors. To gain mechanistic insights, we isolated Mesp1-lineage cells from developing embryos and performed single-cell RNA-sequencing. Our comprehensive single cell transcriptome analysis revealed that Wls deletion dysregulates developmental trajectories of both anterior and posterior SHF cells, marked by impaired proliferation and premature differentiation. Together, these results demonstrate a critical role of local precardiac mesodermal Wnts in SHF fate decision, providing fundamental insights into understanding heart field development and chamber formation.

**Significance Statement:** There is significant interest in understanding the mechanisms underlying heart formation to develop treatments and cures for patients suffering from congenital heart disease. In particular, we were interested in the intricacies of first (FHF) and second heart field (SHF) development, as many congenital heart defects present with heart field-specific etiologies. Here, we uncovered a novel relationship between specified cardiac progenitor cells and second heart field progenitors. Through genetic manipulation of Wnt secretion in developing mouse embryos, we identified a population of cardiac progenitor cells that acts as a local source of Wnts which are necessary for proper SHF development. Our single cell transcriptomic analysis of developing anterior mesoderm showed cardiac progenitor-secreted Wnts function through regulation of differentiation and proliferation among SHF progenitors. Thus, this study provides insight into the source and timing of Wnts required for SHF development, and points to the crucial role of co-developing cell populations in heart development.

---

The heart develops in a synchronized sequence of proliferation and differentiation of cardiac progenitor cells (CPCs) from two anatomically distinct pools of cells, the first and second heart fields (FHF and SHF) (1, 2). Congenital heart defects (CHDs) arise upon dysregulation of these processes, many of which are restricted to derivatives of the FHF or SHF (3). Thus, there is great importance in understanding the crucial steps of how CPCs from each heart field (HF) develop, including identification of the signaling molecules responsible, for better characterization of CHDs.

The patterning and construction of the embryo is controlled by stage and context-specific morphogenic signaling (4). Mesodermal cells form during gastrulation, during which Mesp1^+^ cells migrate anterolaterally across the embryo (5), and are further specified into various cell lineages (6–8), including the FHF and SHF (9). Wnt signaling, in particular, plays an important role in heart development and acts in a stage-specific manner (10–12). Wnt signals are necessary for formation of SHF progenitors from Mesp1^+^ cells (13), and play a very important role in regulation of proliferation and differentiation of SHF CPCs (14–17). Later in development, non-canonical Wnts function to oppose canonical Wnts in SHF (18–20). However, the source of such Wnts regulating heart development remains unknown. Identification of the source will be crucial in understanding the crosstalk between co-developing populations of cells, which will ultimately provide insight into the etiology of various CHDs.

To better understand how the interplay of morphogenic signals mediate HF development, we recently developed a precardiac organoid system which harbors fluorescent reporters of marker genes for each HF (21, 22). In our organoid system, we verified the crucial role of the Wnt signaling pathway in SHF development; however, the source of such Wnts remains elusive. Through bulk transcriptomic analysis of each HF, we found relative upregulation of genes encoding Wnt ligands in the FHF and corresponding upregulation of Wnt receptor and readout genes in the SHF (21). These results point to a potential relationship between the two HFs during development, but a conclusive link has yet to be defined.

To address this, we used an *in vivo* conditional knockout of the gene encoding Wntless (Wls), a protein necessary for Wnt secretion. With this model, we suppressed Wnt secretion from populations of cells and characterized downstream phenotypes. Upon knockout of Wls in anterior mesoderm through Mesp1-Cre (9), heart development was disrupted, with knockout embryos forming a single lone ventricle-like structure. We further characterized this defect as malformation of the SHF, with the ventricle showing FHF features. Subsequent deletion in restricted lineages provided evidence that a CPC population is responsible for secreting Wnts for SHF development. Consistently, single-cell RNA-sequencing (scRNA-seq) revealed that Wnts are enriched in the FHF, and blocking Wnt secretion decreases proliferation of SHF cells, leading to their premature differentiation. Through this study, we provide a basis for SHF cell fate decision - proliferation vs. differentiation - auto-regulated by CPCs.

## Results and Discussion

### Blocking Wnt secretion in anterior mesoderm results in single ventricle phenotype

Wnt signaling is required for SHF development (13); however, the source of Wnts has not yet been identified. While others have perturbed Wnt signaling in Wnt-receiving cells through manipulating *β*-catenin levels, the obligatory transcriptional effector of Wnt signaling, we instead focused on eliminating the ability of cells to secrete Wnts. To suppress Wnt secretion, we used a mouse strain harboring loxP sites flanking the gene encoding Wls (23), a protein required for export of Wnt ligands (24–26). This allowed us to study which types of Wnt-secreting cells affect heart development **(Figure 1A)**.

**Fig. 1.**
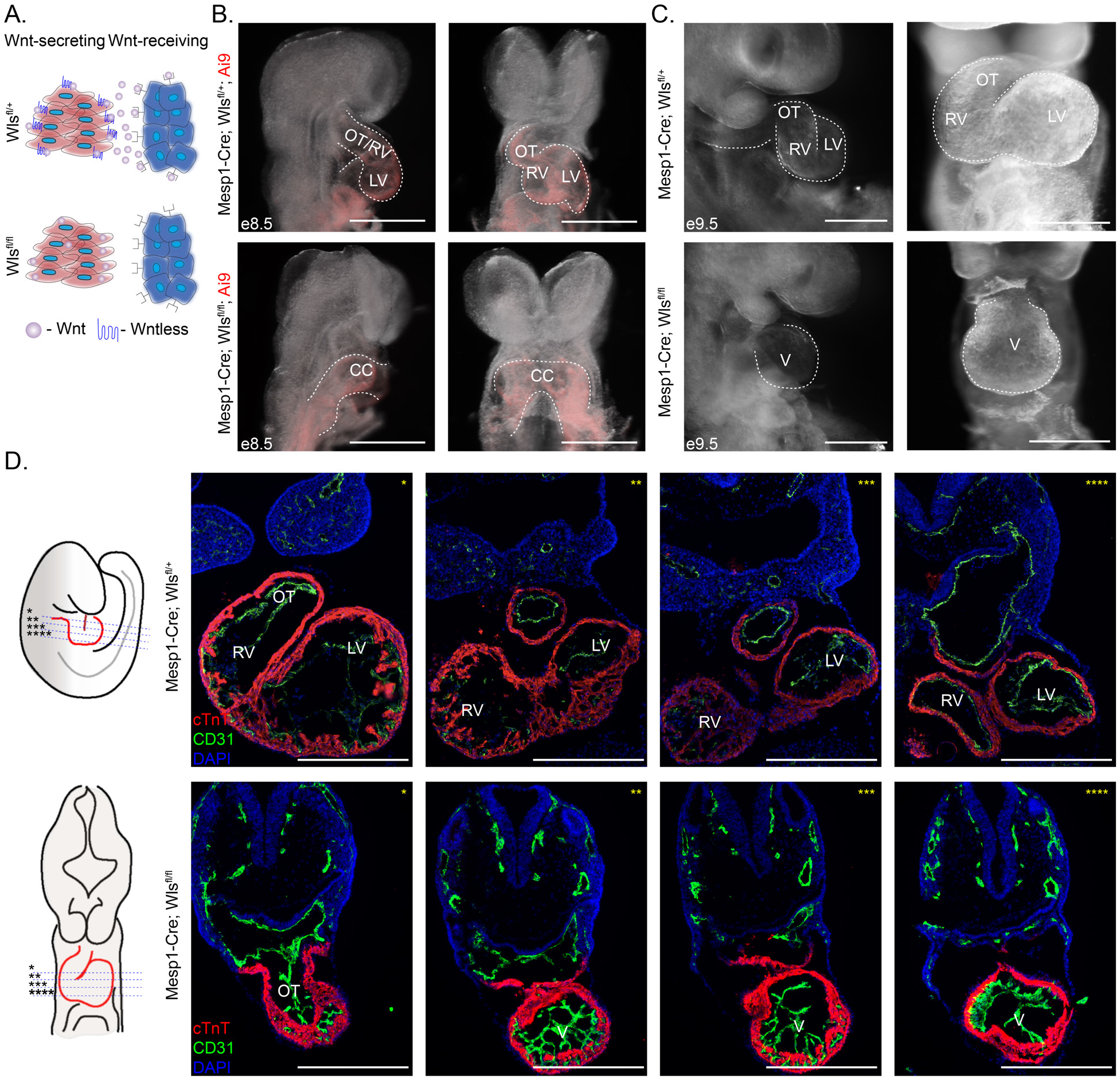
Knockout of Wntless in anterior mesoderm leads to single chamber phenotype. **A.** Schematic detailing conditional knockout of the gene *Wls*. **B.** Side and front images of e8.5 Mesp1-Cre; Wls^*fl/+*^ (control), and Mesp1-Cre; Wls^*fl/fl*^ (knockout) embryos. Ai9 expression marks Mesp1 derivatives. **C.** Side and front images of e9.5 control and knockout embryos. **D.** Schematic illustrating transverse sectioning from anterior to posterior of e9.5 hearts. (CC = Cardiac Crescent, OT = Outflow Tract, RV = Right Ventricle, LV = Left Ventricle, V = Ventricle). White scale bars indicate 500 *μ*m.

We recently found that *Wnt* genes are enriched in FHF progenitors but Wnt activity is higher in SHF progenitors (21). This led us to test if the FHF influences SHF development. As there are no validated Cre drivers that induce recombination with high efficiency in the early FHF, we first deleted Wls in early anterior mesoderm using Mesp1-Cre (9) and traced Mesp1^+^ cell lineage with Ai9. Multiple groups have suggested HF development begins shortly after the onset of Mesp1 expression (6–8). Interestingly, upon knockout of Wls, we found a heart defect starting from embryonic day (e)8.5 **(Figure 1B)**. Control embryos formed the primitive outflow tract/right ventricle (OT/RV) and left ventricles (LV), expected at this developmental stage. However, the cardiac crescent - a structure which is transiently present from e6.0-e8.0 - persisted in knockout embryos at e8.5. Embryo size was found not significantly affected at this stage **(Figure S1A)**. The heart defect became more pronounced by e9.5, where, in comparison to expected LV, RV, and OT formation in control embryos, knockout embryos exhibit a single ventricle-like structure **(Figure 1C)**. Knockout embryos became smaller than control embryos from e9.5 **(Figure S1B-C)**, likely due to the defective heart. Through transverse sectioning, we verified the lone ventricle-like structure throughout the chamber, with no ventricle chamber septation present **(Figure 1D)**.

### SHF structures are malformed without Wnt signals from the anterior mesoderm

The presence of the cardiac crescent (FHF) in knockout embryos suggests that the single ventricle phenotype may be due to dysregulation of SHF development. To test this, we utilized an Hcn4-GFP reporter mouse strain (27, 28) to visualize FHF progenitors. As expected, GFP+ cells were localized in the LV of control embryos. However, knockout embryos showed an expansion of the GFP expressing domain across the entire ventricle-like structure **(Figure 2A, Figure S1D)**. Consistently, immunostaining showed that the entire ventricle was positive for the LV marker Tbx5 in knockout embryos while only the LV was positive for Tbx5 in control embryos **(Figure 2B)**, supportive of a lack of SHF derivatives. However, SHF progenitors were still present where the single chamber is attached to pharyngeal arches in knockout embryos **(Figure S1E)** though there is a notable decrease in Mesp1 lineage cells in the second pharyngeal arches (PA2) that contain SHF cells (29) **(Figure 2C)**. These results suggest that mesodermal *Wnts* are required for SHF development, but not for specification.

**Fig. 2.**
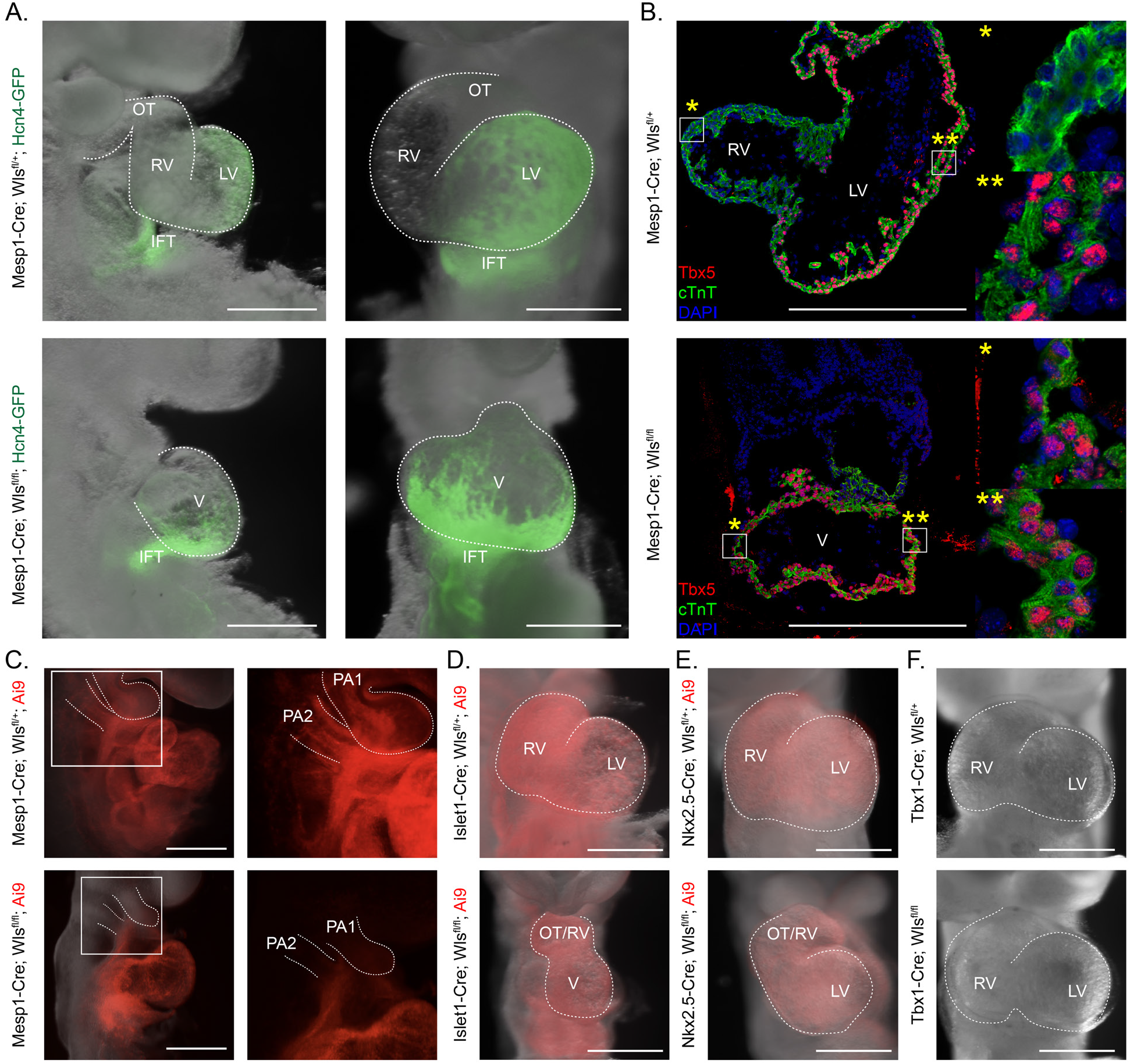
Lone ventricle formed is a FHF derivative and is a result of Wnt signaling from non-SHF cardiac mesoderm. **A.** Fluorescent overlay images of Hcn4-GFP reporter in Mesp1-Cre; Wls^*fl/+*^ (control) and Mesp1-Cre; Wls^*fl/fl*^ (knockout) embryos. **B.** Immunostaining of control and knockout sections of embryo hearts for cTnt (green), a cardiomyocyte marker, and Tbx5 (red), a left ventricle marker, with emphases on the right (*) and left (**) sides of the resulting chambers. **C.** Immunofluorescence of the Mesp1^+^ lineages (Ai9) in control and knockout embryos, with focus on the pharyngeal arch regions. **D.** Front images of Islet-cre control and knockout embryos. Ai9 marks Islet1 derivatives. **E.** Front images of Nkx2.5-Cre control and knockout embryos. Ai9 marks Nkx2.5 derivatives. **G.** Front images of Tbx1-Cre control and knockout embryos. (OT = Outflow Tract, IFT = Inflow Tract, RV = Right Ventricle, LV = Left Ventricle, V = Ventricle, PA1 = 1st Pharyngeal Arch, PA2 = 2nd Pharyngeal Arch). White scale bars indicate 500 *μ*m.

### CPCs Wnts are required for SHF development

To determine the source of mesodermal Wnts more specifically, we leveraged multiple precardiac Cre drivers with shared expression domains to conditionally knockout Wls. We first used Islet1-Cre mice, as its Cre expression begins shortly after gastrulation in FHF progenitors before marking the SHF (21, 30, 31). We chose this particular Cre-driver as it is activated in both heart fields and expressed early in heart development, which we confirmed by observing RFP expression in both LV and RV in Cre-recombined control embryos **(Figure 2D)**. Interestingly, we observed a similar single ventricle phenotype in Islet1-Cre; Wls knockout embryos **(Figure 2D)**, albeit with variable penetrance. These results suggest that the source of Wnts is from a common population of cells derived from both Mesp1^+^ and Islet1+ cells - the FHF and SHF. Supporting this interpretation, we observed a hypoplastic OT/RV when deleting Wls with Nkx2.5-Cre mice **(Figure 2E)**. Nkx2.5 is expressed in CMs of both HFs, at a slightly later stage of development than Mesp1 and Islet1, which potentially explains the decreased penetrance and severity of the SHF defect. To further test which heart field is responsible for the phenotype, we blocked Wnt secretion in the SHF using Tbx1-Cre mice. No heart defect was observed in Tbx1-Cre; Wls knockout embryos **(Figure 2F)**. This may indicate that SHF Wnts are dispensable for SHF development. Together, these results support that HF cells provide Wnts for SHF development. Further study to more specifically identify the responsible cell population is needed, as it is possible that the observed Mesp1/Islet1/Nkx2.5 phenotypes may be resulting from independent mechanisms. Nevertheless, our results provide the first evidence of a source of Wnts essential for SHF development.

### scRNA-seq identifies perturbation of mesodermal development in Wls knockout

To understand the nature of the observed defect at the transcriptional level, we isolated Mesp1^+^ cells from *Mesp1-Cre; Ai9; Wls*^*fl/+*^ *and Wls*^*fl/fl*^ mice at three developmental timepoints (e8.0, e8.5, and e9.5) and performed scRNA-seq **(Figure 3A)**. As separating control and knockout cells into separate capture runs can lead to potential downstream batch effects (32), we multiplexed and pooled samples using lipid-tagged indices through MULTI-seq (33). We subsequently performed fluorescence-activated cell sorting to isolate RFP+ cells, and generated sequencing libraries through the 10x Chromium platform. We performed sequencing across 3 runs, and combined and integrated the data using *scTransform* normalization followed by anchoring gene integration as implemented in Seurat (32, 34, 35). Our final dataset, after quality control and filtration of red blood cells, was composed of 11,959 cells.

**Fig. 3.**
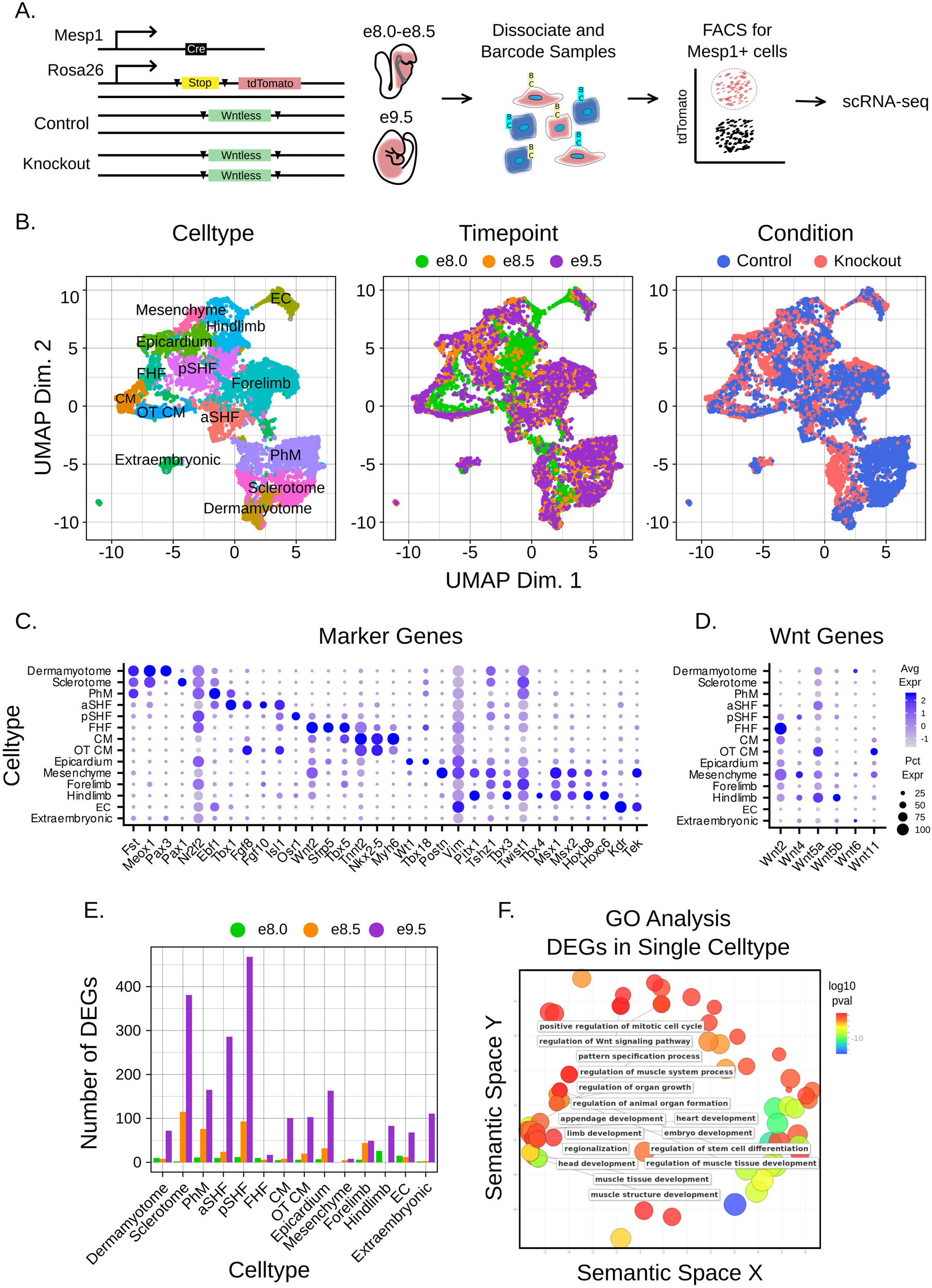
SHF development is dysregulated through decreased proliferation without anterior mesoderm Wnts. **A.** Experimental design for single cell RNA-seq experiment. **B.** UMAP clustering based on cell type, timepoint, and condition. **C.** Marker gene expression used to identify cell populations.**D.** Expression of *Wnt* genes across cell populations. **E.** Quantification of differentially expressed genes (DEGs) across timepoints and cell populations. **F.** GO terms from DEGs found uniquely in one cell population.

We performed dimensionality reduction through UMAP and clustering through Seurat (32, 35) **(Figure 3B)**. We subsequently annotated clusters through known marker genes **(Figure 3C)**. Our full cluster identification strategy is detailed in the **Supplementary Methods** section. The dataset contained well-known subpopulations of mesodermal origin, including both HFs (FHF, anterior and posterior SHF [aSHF, pSHF]), somites (dermamyotome and sclerotome), pharyngeal mesoderm (PhM), cardiomyocytes (CMs, including outflow tract [OT] CMs), other cardiac tissues (endothelial/endocardium [EC], epicardium, mesenchyme), other mesodermal populations (forelimb and hindlimb), and extraembryonic mesoderm. We then analyzed Wnts in each population of mesoderm. Based on levels of expression of relevant Wnts, and our Wls loss of function studies, we focused on three populations as putative sources of instructive Wnts - pSHF, FHF, and OT CM - though the OT CMs are known to derive from Tbx1+ cells (21, 36, 37). The FHF highly expressed *Wnt2*, which corroborated our previous findings (21), while the pSHF expressed low/undetectable levels of *Wnts* **(Figure 3D)**. The OT CMs highly expressed *Wnt5a* and *Wnt11*, which are known for their roles in OT development (18–20). This data further supports our hypothesis that the FHF may be a source of instructive Wnts.

Expression levels of *Wls* were abrogated in nearly every knockout tissue (with the exception of the extraembryonic mesoderm), supporting successful knockout of the gene **(Figure S2A)**. Though the level of our sampling does not allow for population-based conclusions based on percentage of cells, we quantified the fraction of cells allocated to each population in control and knockout animals **(Figure S2C)**. While some differences were observed, there was minimal difference between the proportion of aSHF cells in control and knockout, which may suggest that Wls deletion does not influence SHF specification. However, several clusters of knockout cells separated entirely from control cells in multiple populations **(Figure 3B)**, supporting transcriptional changes following Wls knockout. We performed differential gene testing between control and knockout cells and found relatively few transcriptional differences at e8.0 and progressively more changes at e8.5 and e9.5 **(Figure 3E)**. The populations most affected included the sclerotome, PhM, aSHF, pSHF, and epicardium.

To investigate global transcriptional effects, we first identified a conserved pool of 76 genes that were differentially expressed in at least 7 of the 14 populations **(Figure S2B)**. This pool was enriched for genes involved in tissue stress response. In particular, knockout cells showed upregulation of known apoptosis genes **(Figure S2D)**, upregulation of glycolytic genes **(Figure S2E)**, and downregulation of genes involved in mitochondrial and oxidative function **(Figure S2F)**. These results are supportive of a generalized tissue response in Wls knockout, corresponding to the knockout phenotype.

We then asked how Wls deletion affected mesodermal cells outside of generalized response due to secondary effects of having a defective heart. To do this, we explored genes that were differentially expressed in unique clusters of cells **(Fig S2G)**. Notably, this pool of genes was enriched for Gene Ontology terms related to cell cycle, Wnt signaling, and mesodermal organ tissue and development **(Figure 3F)**. This suggested that, outside of the nonspecific stress response in all tissues, Wls deficiency leads to perturbation of tissue development in specific mesodermal processes. The majority of these differentially expressed genes were identified in aSHF, pSHF, and sclerotome **(Figure S2G)**, suggesting these tissues are most perturbed by loss of Wnt signaling.

### Failure of proliferation and premature differentiation underlie aSHF defect

Given the perturbation of SHF cells, we performed trajectory reconstruction of control and knockout cells in the aSHF using Monocle 2 (38). Monocle 2 reconstructed a branched trajectory in which control and knockout cells are largely indistinguishable early at e8.0, before reaching separated states at later timepoints **(Figure 4A)**. We subsequently used tradeSeq (39) to identify genes differentially expressed across the control and knockout paths at the branch point, focusing on genes with differential expression at the end states. We identified 753 genes, which we classified into four clusters based on gene expression dynamics **(Figure 4B)**. In general, GO terms relevant to heart development were enriched in upregulated gene clusters while GO terms related to cell cycle were enriched in downregulated clusters **(Figure 4B)**. We focused on specific sets of candidate genes to verify trends of expression. First, we looked at the cell cycle progression genes *Ccnd1*, *Ccne1*, *Cdc6*, *Cdc20*, and *Nuf2*, and observed that they were indeed downregulated, while the cell cycle inhibitor *Ccng2* was upregulated **(Figure 4C)**, pointing to potential impaired proliferation. Given that proper SHF development requires maintenance of proliferation and delayed differentiation, we found the accompanied upregulation of aSHF and cardiomyogenesis genes particularly intriguing. Specifically, upregulation of the cardiomyocyte genes *Nkx2-5*, *Tgfb1*, and *Smad6* (important for inhibition of BMP signaling for cardiomyogenesis) (40, 41), *Ttn* and *Nebl* (part of the sarcomeric apparatus), and *Tbx3* **(Figure 4C)** are consistent with this hypothesis. The aSHF genes *Fgf8* /*10*, *Isl1*, *Mef2c* and *Hand2* were all upregulated while *Nr2f1* was downregulated, though their trends were more complicated **(Figure 4C)**. The late upregulation of aSHF genes that regulate proliferation (*Fgf8* /*10*) was particularly interesting, and points to a potential dysregulation of cell identity after premature differentiation. This suggests that SHF progenitors are pre-differentiated, but may remain in a niche (rather than migrate to the heart), causing further dysregulation. We verified this *in vivo* using EDU in the pan-cardiac deletion of Wls in the Islet1 lineage. Knockout embryos showed significant decrease in cycling cells in pharyngeal arches, where aSHF cells undergo proliferation (29) **(Figure S1G)**. Taken together, these results point to a premature differentiation of aSHF progenitors and decrease in cell proliferation as a key mechanism for the knockout heart phenotype.

**Fig. 4.**
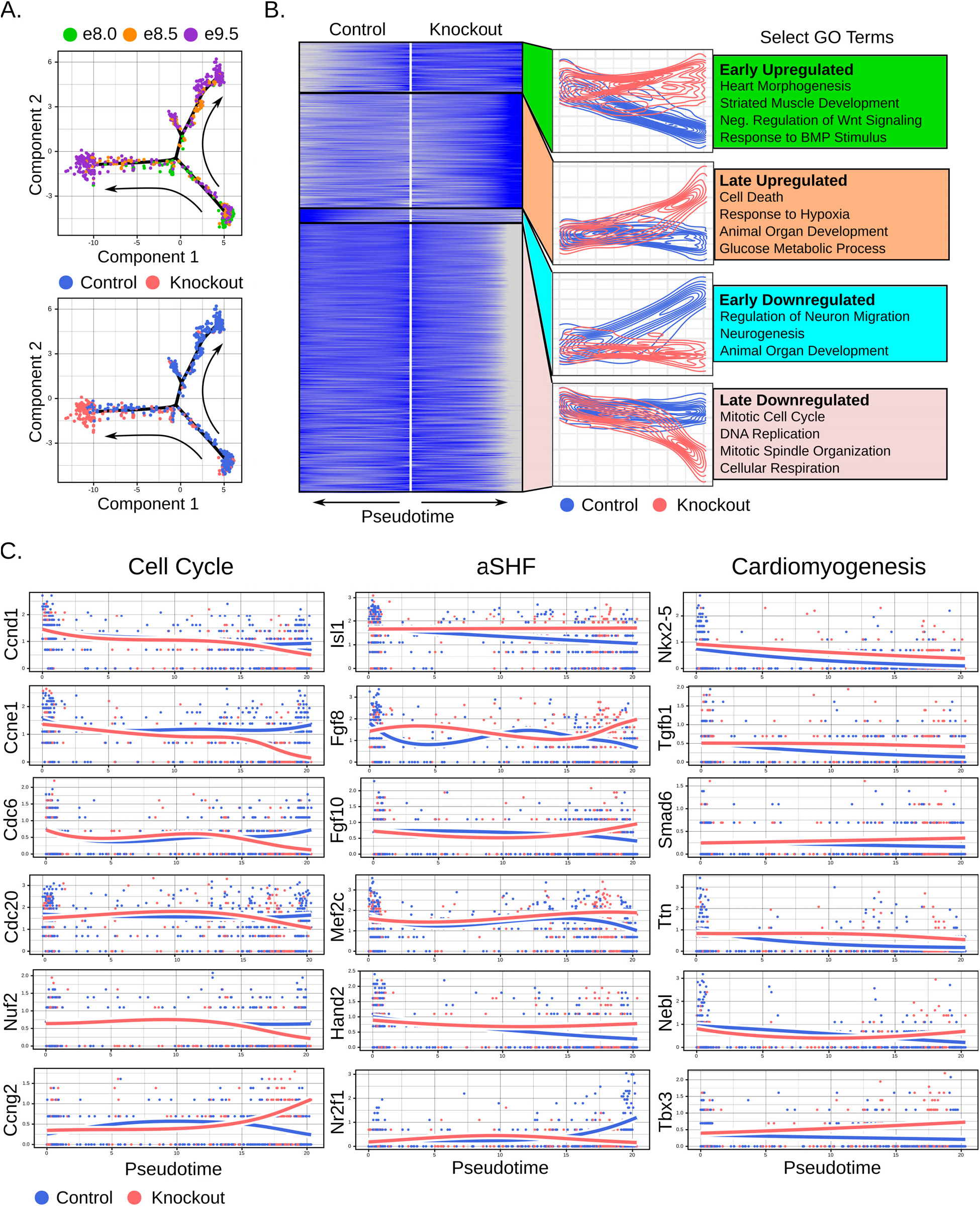
aSHF development is impaired by premature differentiation and reduced proliferation in Wls knockout embryos. **A.** Developmental trajectories from aSHF of control and knockout embryos. **B.** Heatmap identifying four clusters of differential gene trends. **C.** Gene expression trends across trajectory of relevant cell cycle, aSHF, and cardiomyogenic genes.

We then performed a similar analysis with pSHF cells, and again observed a branched trajectory with early time points broadly aligning and later timepoints clustering on different branches **(Figure S3A)**. In addition to heart morphogenesis and cell cycle gene dysregulation in knockout pSHF cells, the significant upregulation of apoptosis genes in knockout cells suggests that Wnt signals are responsible for maintaining pSHF cell survival **(Figure S3B-C)**. The differing results among aSHF and pSHF supports the context-dependent nature of Wnt signaling, and interestingly even among the SHF during development.

In the present study, we demonstrate a novel auto-regulatory role for the precardiac mesoderm in instructing the SHF development by regulation of proliferation and differentiation of SHF progenitors. While further study would be necessary to pinpoint the temporospatial dynamics of Wnts, our study provides the first evidence that cardiac progenitors secrete Wnts to determine SHF cell fate. Organoid models (42) may help with greater control of conditions and access for sampling in the future.

## Materials and Methods

All methods, including wet lab and computational methods, can be found in the Supplementary Information. Raw data for the Wls knockout scRNA-seq experiments can be found on GEO at GSE165300. Code to generate figures in this manuscript as well as the counts tables for the datasets analyzed in this manuscript can be found on Github at https://github.com/skannan4/wls, and further files can be found on Synapse (https://www.synapse.org/#!Synapse:syn24200678/files/).

## ACKNOWLEDGMENTS

We thank Linda Orzolek and the Johns Hopkins Transcriptomics and Deep Sequencing core for 10x library prep and sequencing. We also thank Dr. George McNamara for assistance and training in confocal microscopy. MULTI-seq reagents were kindly provided by the Gartner lab at UCSF, with assistance graciously provided by Chris McGinnis. Additionally, we thank members of the Dr. Chulan Kwon lab for helpful discussion. This work was supported by grants from National Institutes of Health, American Heart Association, Maryland Stem Cell Research Fund, and the Department of Defense.

## Supplementary Information for

### Supporting Information Text

The purpose of our supplementary materials section is to provide detailed information about our study that was not included in the main manuscript text. We aspire to standards of data reproducibility and availability. To this end, all of the sequencing data for this study can be found on GEO with accession number GSE165300. Additionally, the code to reproduce our findings is available on Github at https://github.com/skannan4/wls. Lastly, we have made an R workspace available on Synapse (https://www.synapse.org/#!Synapse:syn24200678/files/) that contains many of the data tables pre-loaded for our analysis. If there are other materials that could facilitate re-analysis or exploration, we would be glad to provide them on inquiry.

### Supplementary Methods

#### Animal Work

Mesp1-Cre (1), Tbx1-Cre (2), Islet1-Cre (3), Nkx2.5-Cre (4), Hcn4-GFP (5, 6), Wntless loxp (stock no. 027484, Jackson Laboratory, (7)), and Ai9 (stock no. 007909, Jackson Laboratory, (8)) mouse strains were utilized for *in vivo* experiments. Embryos were harvested from embryonic day (e)8.0-10.5 for further analysis. Each embryo was genotyped using tissue from individual yolk sacs. Embryos intended for cryo-sectioning and immunofluorescence were cleaned, fixed in 4% PFA for two hours, and stored in 30% sucrose overnight. Embryos were then embedded in OCT and flash frozen before sectioning. Antibodies used in this project were Tbx5 (Atlas Antibodies; HPA008786), cTnt (Thermo Fisher; MS-295-P1), CD31 (BD Biosciences; 553371), Islet1 (DSHB; 39.3F7), Nkx2-5 (SCBT; sc-8697). Stained sections were imaged on a Keyence BZ-X710 microscope. Whole mount EDU experiments were completed as described previously (9). Embryos intended for transcriptomic analysis were dissected, staged, and stored on ice in cold PBS without Ca^2+^ or Mg^2+^. Embryos were dissociated and barcoded using MULTI-seq anchor and primers (10). Subsequently, Mesp1^+^/RFP cells were isolated by FACS using Sony SH800. Cells were then captured for 10x library prep and single cell RNA sequencing. All protocols involving animals followed U.S NIH guidelines and were approved by the ACUC of JHMI.

#### scRNA-seq Library Preparation and Sequencing

We multiplexed samples for sequencing using the MULTI-seq protocol (10). We adapted the protocol described below from the instructions provided to us by the Gartner lab. The following primers/adapters were used:

- Anchor LMO: 5’-TGGAATTCTCGGGTGCCAAGGgtaacgatccagctgtcact-{Lipid}-3’
- Co-Anchor LMO: 5’-{Lipid}-AGTGACAGCTGGATCGTTAC-3’
- Barcode Oligo: 5’-CCTTGGCACCCGAGAATTCCANNNNNNNNA30-3’
- MULTI-seq Primer: 5’-CTTGGCACCCGAGAATTCC-3’
- TruSeq RPIX: 5’-CAAGCAGAAGACGGCATACGAGATNNNNNNGTGACTGGAGT
- TCCTTGGCACCCGAGAATTCCA-3’
- Universal I5: 5’-AATGATACGGCGACCACCGAGATCTACACTCTTTCCCTACACGACGCTCTTCCGATCT-3’

The anchor and co-anchor LMOs were kindly provided to us by the Gartner lab; the barcode oligos, MULTI-seq primer, and TruSeq RPIX were ordered from Integrated DNA Technologies using barcodes provided by the Gartner lab; and the Universal i5 was part of the 10x Chromium 3’ v3 Reagent kit.

Briefly, after sorting, cells were incubated with a 10x anchor:barcode solution for 5 minutes on ice, followed by incubation with 10x co-anchor solution for 5 minutes. We subsequently added 1% BSA in PBS and washed 2-3 times. We used the 10x Chromium 3’ v3 workflow to encapsulate and capture cells, reverse transcribe mRNA, and purify cDNA. To capture the barcode sequences, we performed the 10x cDNA amplification reaction, but with addition of the MULTI-seq primer. Cleanup with 0.6x SPRI beads enabled separation of endogenous transcript cDNA and barcodes - endogenous transcripts remained bound to the beads while barcodes were eluted in the supernatant. We then separately processed the endogenous cDNA and barcodes using the 10x workflow. For the experiments sequenced on 10/21/19, sequencing of the endogenous transcripts was done on a NextSeq500 high output lane using a 28/8/91 bp design for R1/i7/R2. Sequencing of the barcodes was done a NextSeq500 mid output lane using a 28/8/8 design for R1/i7/R2. For the experiments sequenced on 03/04/20, sequencing for both endogenous and barcode transcripts was done on a NovaSeq6000 S1 flow cell, using a 28/8/91 bp design; the barcode transcripts were subsequently trimmed using the BBMap/BBTools suite (11).

#### Mapping and Demultiplexing

Endogenous cDNA reads were mapped using Kallisto|Bustools (0.46.1) (12). We removed poor quality barcodes and demultiplexed the samples using deMULTIplex (1.0.2) (10), with minor modifications as described in our code. We removed all cells classified as “Negative” or “Doublet”. Our final analysis yielded the following samples:

- Run 1 (10/21/19): 1 e8.5 KO, 2 e8.5 WT, 1 e9.5 KO, 2 e9.5 WT
- Run 2 (10/21/19): 1 e7.5 WT, 2 e9.5 KO, 2 e9.5 WT
- Run 3 (03/04/20): 3 e8.0 KO, 3 e8.0 WT, 3 e9.5 KO, 2 e9.5 WT

We performed further quality control by removing low quality cells and putative doublets using the following selection criteria - 2500 < genes < 9000; mitochondrial percentage < 22%; total UMIs < 62500.

#### Dataset Integration, Dimensionality Reduction, and Clustering

Integration of the three runs was performed using the *SCTransform* workflow in the Seurat package (3.1.4) (13) as described by the vignette provided by the authors. In particular, we selected 3000 features for integration. Following integration, we found little evidence of batch-specific clustering, suggesting successful integration of the three runs. Dimensionality reduction via UMAP and clustering (using the Louvain algorithm) were performed in Seurat. We selected 15 principal components for clustering, as an elbow plot indicated that these were sufficient for capturing the majority of variation in the dataset; moreover, we found that increasing the number of principal components led to abiological clustering. As discussed below, we overclustered by using a resolution of 2.0 and subsequently merged similar clusters.

#### Clustering and Annotation Strategy

There is no established method for identifying the optimal number or size of clusters for an scRNA-seq experiment, though various metrics and strategies have been proposed (14). Indeed, the appropriate clustering approach will vary from experiment to experiment based on the goals and desired resolution. Our strategy here was to purposely overcluster (e.g. generate more clusters than biologically expected) and then manually merge and label clusters based on expression of marker genes of interest.

Here, we discuss the markers we selected for identification of clusters. We utilized a set of known canonical markers to identify each cluster of cells. Clusters that were labeled as the same cell identity were merged. Our initial clustering was done to remove RBCs. We then determined somitic mesoderm (sclerotome and dermomyotome), through high expression of the gene *Meox1* (15). We were able to differentiate between sclerotome and dermomyotome through expression of *Pax1* (16–19) (sclerotome) and *Pax3* (16, 20, 21) (dermomyotome). Next, we utilized a recent very thorough single cell analysis of heart development (22), as well as other studies, to identify cardio-pharyngeal clusters through published marker genes. We determined our pharyngeal mesoderm (PhM) through co-expression of *Fst*, *Ebf1* and *Tbx1*. We determined anterior SHF (aSHF) identity through co-expression of *Fgf8* /*10*, *Isl1*, *Tbx1*, and *Sema3c*. Posterior SHF (pSHF) identity was determined through upregulation of *Osr1*, *Hoxb1*, and *Foxf1*, with mild *Isl1* expression. The FHF was identified as a cluster that was mostly present at e8.0, and expressed FHF markers *Wnt2*, *Sfrp5*, *Tbx5*, and mild expression of the cardiomyocyte markers *Tnnt2* and *Nkx2-5*. Cardiomyocytes were identified by cardiomyocyte markers including *Tnnt2*, *Nkx2-5*, and *Myh6*. OT CMs were identified specifically by co-expression of CM markers with *Fgf8* and *Isl1*. The epicardium was identified through *Wt1* and *Tbx18* co-expression (23, 24). Mesenchymal cells were identified through co-expression of *Postn* (25), *Vim* (26), *Pitx1*, *Twist*, and *Tek*. Forelimb identity was established through co-expression of *Fgf10*, *Tbx5* (27, 28), *Tshz2*, and *Lmx1b* (29). We differentiated between forelimb and hindlimb through *Tbx4* and *Tbx5* expression (forelimb expressing *Tbx5* and hindlimb expressing *Tbx4*) (30), in addition to expression of *Pitx1* (31), *Hoxb8* (32), and *Hoxc6* (33). We identified the endocardium/endothelial (EC) identity through co-expression of *Kdr* and *Tek* (34). Remaining cells which clustered separately from other cell types and showed no discernable gene expression patterns were identified as extraembryonic in origin.

#### Differential Gene Expression Testing and Analysis

Differential expression testing (as in **Figure 3F**) was performed by comparing control to knockout cells in each population at each timepoint using the Wilcoxon rank sum test as implemented as Seurat. For a comparison to be performed, we required at least 10 cells to be present in both the control and knockout. A minimum expression of 25% of cells was used as a cutoff for testing genes. We subsequently selected genes with Bonferroni correction-adjusted p-value < 0.05. We performed Gene Ontology Overrepresentation Analysis by inputting differentially expressed genes into the the online Gene Ontology Resource (35, 36). Summary and visualization of GO terms was done by selecting the top 150 GO terms by fold enrichment and inputting to REViGO (37). For **Figure 3G**, we selected genes that were differentially expressed in only one tissue at e9.5, while for **Figure S2F**, we selected genes that were differentially expressed in at least 7 tissues.

#### Trajectory Reconstruction and Analysis

Trajectory reconstruction was performed using Monocle 2 (2.12.0) (38); while we initially tested Monocle 3 and Slingshot, we found that Monocle 2 offered the most flexibility and yielded trajectories that most matched the expected biology. We used a semi-supervised approach to generating trajectories. We first selected the 8000 most variable genes; these genes were provided as input to differential gene expression testing between the control and knockout cells at e9.5. We subsequently selected the 3000 most differentially expressed genes to construct the trajectory. This approach potentially biases to differences between control and knockout at the expense of other biological variation. We favored this approach because our specific goal was to understand the deviation in the states of the control and knockout cells. However, we found that an unbiased approach (dpFeature, as implemented in Monocle 2) yielded similar trajectories, suggesting that the control vs knockout differences are the most notable biological variation in our tested cells. We performed differential gene expression testing across the branches by using tradeSeq (1.3.15) (39). As the pseudotimes from Monocle 2 may be somewhat arbitrary across branches, we stretch branches such that they had the same overall length and end pseudotime. We additionally pruned small side branches that were likely artefactual. We fit generalized additive models to genes expressed in at least 20% of cells across the branches, and identified genes differentially expressed across the end states using the diffEndTest function in tradeSeq. The p-values reported by tradeSeq are to be interpreted with some caution; however, simply to specify a threshold for further analysis, we focused on genes with Benjamini-Hochberg-corrected p-values < 0.05 and log2(Fold Change) >= 0.8. We clustered gene trends using complete-linkage hierarchical clustering as implemented in the pheatmap package (1.0.12).

---

**Fig. S1.**
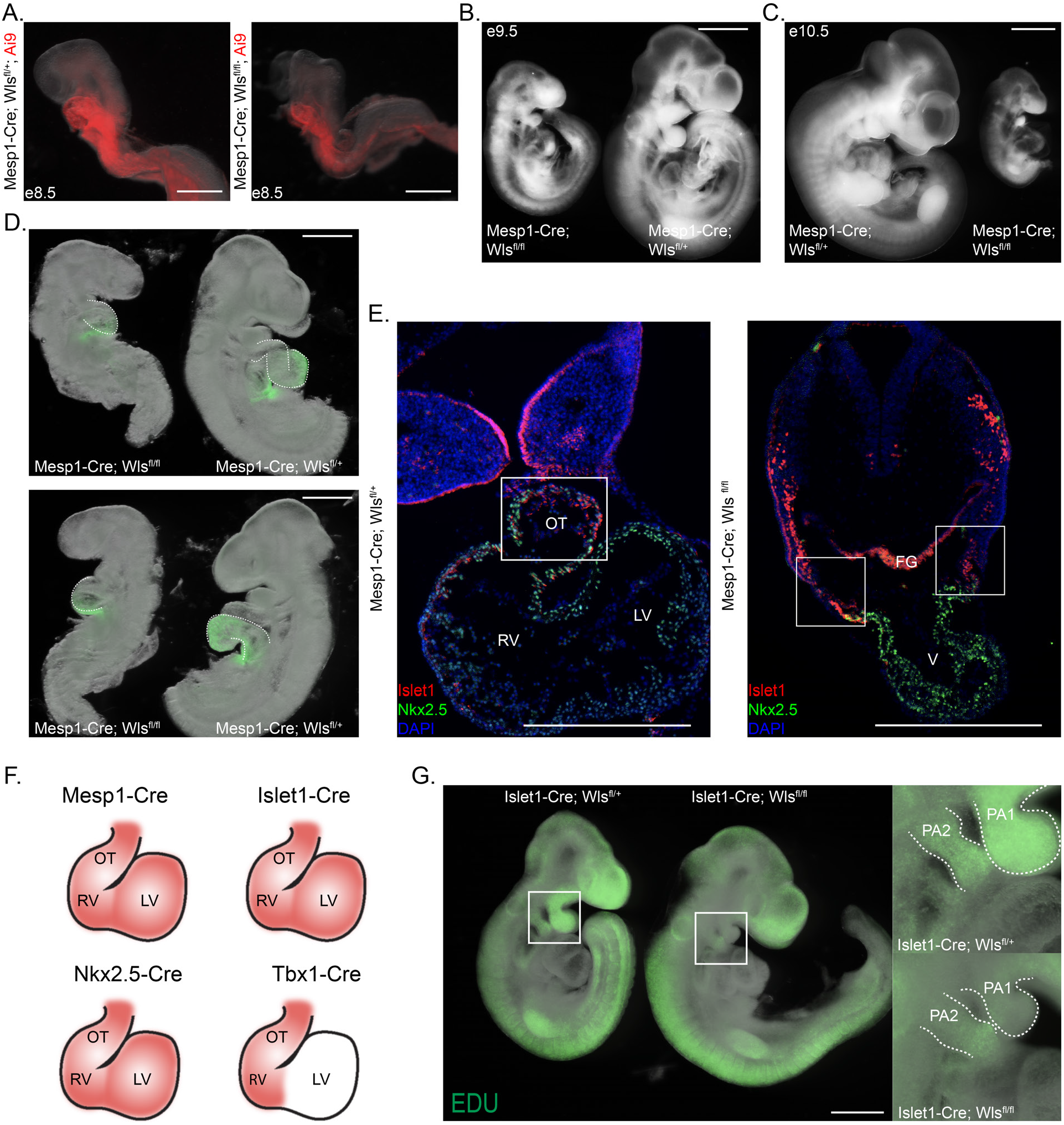
*In vivo* analysis supports global and SHF-specific phenotypes in knockout embryos. **A.** Left side images of full Mesp1-Cre control and knockout e8.5 embryos. **B.** Right side images of full Mesp1-Cre control and knockout e9.5 embryos. **C.** Right side images of full Mesp1-Cre control and knockout e10.5 embryos. **D.** Right and left side images of full Mesp1-Cre control and knockout e9.5 embryos with Hcn4-GFP labeling. **E.** Transverse sections of E9.5 control and knockout embryos showing the presence of Islet1 progenitors. **F.** Schematic detailing expression domains of Mesp1, Islet1, Nkx2.5, and Tbx1 lineages. **G.** EDU stain of Islet1 control and knockout E9.5 embryos with focus on PA1 and PA2 regions. (OT = Outflow Tract, RV= Right Ventricle, LV = Left Ventricle, FG = Foregut Endoderm, V = Ventricle, PA1 = 1st Pharyngeal Arch, PA2 = 2nd Pharyngeal Arch). White scale bars indicate 500 *μ*m.

**Fig. S2.**
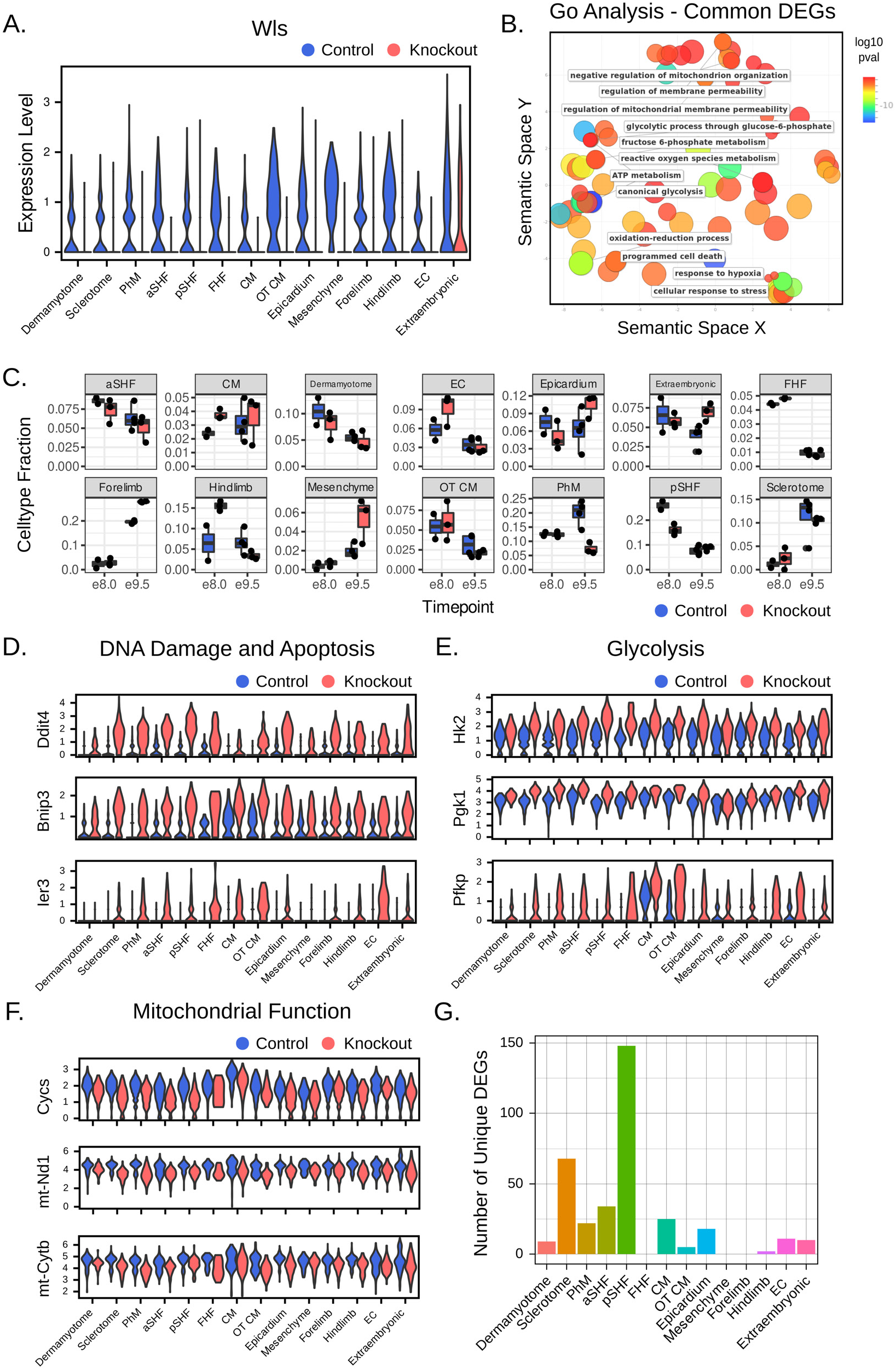
scRNA-seq identifies disruption of heart development in Wls knockout. **A.** Violin plots showing successful knockout of Wls in Mesp1^+^ cell populations. **B.** GO terms enriched in 7/14 cell populations. **C.** Quantification of cell populations in control and knockout embryos per timepoint. **D.** Gene expression across cell populations of control and knockout embryos related to DNA damage and apoptosis. **E.** Gene expression across cell populations of control and knockout embryos related to glycolysis. **F.** Gene expression across cell populations of control and knockout embryos related to mitochondrial function. **G.** Quantification of total unique DEGs per cell population between control and knockout embryos.

**Fig. S3.**
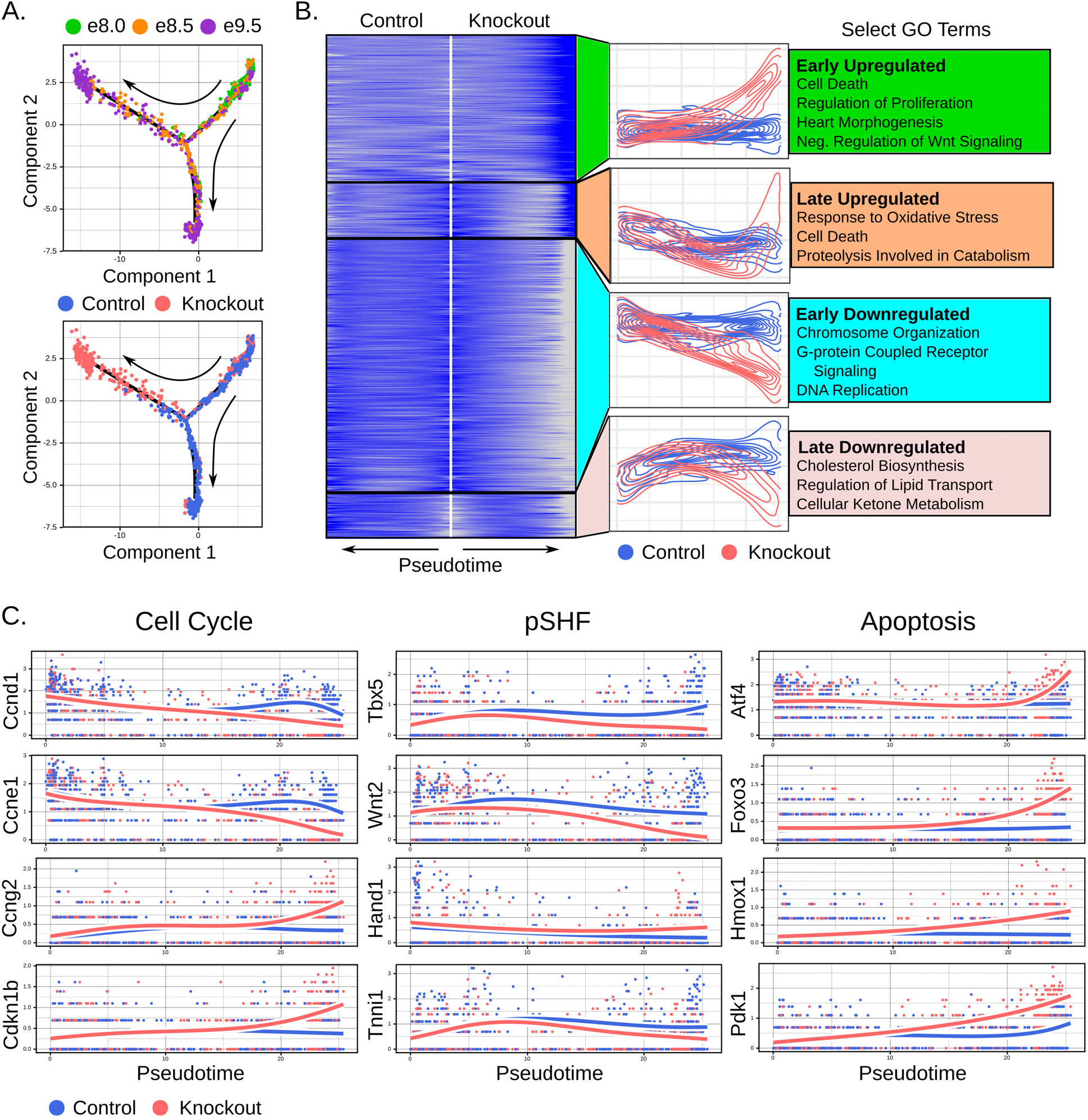
pSHF survival and proliferation is impaired in Wls knockout embryos. **A.** Developmental trajectories from pSHF of control and knockout embryos. **B.** Heatmap identifying four clusters of differential gene trends. **C.** Gene expression trends across trajectory of relevant cell cycle, pSHF, and apoptosis genes.

## References

1. M Buckingham, S Meilhac, S Zaffran, Building the mammalian heart from two sources of myocardial cells. Nat. Rev. Genet. 6, 826–835 (2005).

2. SM Meilhac, ME Buckingham, The deployment of cell lineages that form the mammalian heart (2018).

3. X Liu, et al., The complex genetics of hypoplastic left heart syndrome. Nat. Genet. 49, 1152–1159 (2017).

4. FX Galdos, et al., Cardiac Regeneration: Lessons from Development (2017).

5. S Kitajima, A Takagi, T Inoue, Y Saga, MesP1 and MesP2 are essential for the development of cardiac mesoderm. Development 127(2000).

6. WP Devine, JD Wythe, M George, K Koshiba-Takeuchi, BG Bruneau, Early patterning and specification of cardiac progenitors in gastrulating mesoderm. eLife 3(2014).

7. F Lescroart, et al., Early lineage restriction in temporally distinct populations of Mesp1 progenitors during mammalian heart development. Nat. Cell Biol. 16, 829–840 (2014).

8. F Lescroart, et al., Defining the earliest step of cardiovascular lineage segregation by single-cell RNA-seq. Science 359, 1177–1181 (2018).

9. Y Saga, et al., MesP1 is expressed in the heart precursor cells and required for the formation of a single heart tube. Development 126, 3437–3447 (1999).

10. C Kwon, KR Cordes, D Srivastava, Wnt/*β*-Catenin signaling acts at multiple developmental stages to promote mammalian cardiogenesis. Cell Cycle (2008).

11. L Lin, et al., *β*-Catenin directly regulates Islet1 expression in cardiovascular progenitors and is required for multiple aspects of cardiogenesis. Proc. Natl. Acad. Sci. United States Am. 104, 9313–9318 (2007).

12. AT Naito, et al., Developmental stage-specific biphasic roles of Wnt/-catenin signaling in cardiomyogenesis and hematopoiesis. Proc. Natl. Acad. Sci. United States Am. (2006).

13. A Klaus, Y Saga, MM Taketo, E Tzahor, W Birchmeier, Distinct roles of Wnt/-catenin and Bmp signaling during early cardiogenesis, Technical report (2007).

14. D Ai, et al., Canonical Wnt signaling functions in second heart field to promote right ventricular growth. Proc. Natl. Acad. Sci. United States Am. (2007).

15. ED Cohen, et al., Wnt/*β*-catenin signaling promotes expansion of Isl-1–positive cardiac progenitor cells through regulation of FGF signaling. J. Clin. Investig. 117, 1794–1804 (2007).

16. C Kwon, et al., Canonical Wnt signaling is a positive regulator of mammalian cardiac progenitors. Proc. Natl. Acad. Sci. United States Am. (2007).

17. Y Qyang, et al., The Renewal and Differentiation of Isl1 + Cardiovascular Progenitors Are Controlled by a Wnt/b-Catenin Pathway. Cell Stem Cell (2007).

18. M Abdul-Ghani, et al., Wnt11 Promotes Cardiomyocyte Development by Caspase-Mediated Suppression of Canonical Wnt Signals †. MOLECULAR AND CELLULAR BIOLOGY 31, 163–178 (2011).

19. JA Bisson, B Mills, JCP Helt, TP Zwaka, ED Cohen, Wnt5a and Wnt11 inhibit the canonical Wnt pathway and promote cardiac progenitor development via the Caspase-dependent degradation of AKT. Dev. Biol. 398, 80–96 (2015).

20. ED Cohen, MF Miller, Z Wang, RT Moon, EE Morrisey, Wnt5a and wnt11 are essential for second heart field progenitor development. Development 139, 1931–1940 (2012).

21. P Andersen, et al., Precardiac organoids form two heart fields via Bmp/Wnt signaling. Nat. Commun. 9, 3140 (2018).

22. E Tampakakis, M Miyamoto, C Kwon, In vitro generation of heart field-specific cardiac progenitor cells. J. Vis. Exp. 2019, e59826 (2019).

23. J Fu, HM Ivy Yu, T Maruyama, AJ Mirando, W Hsu, Gpr177/mouse Wntless is essential for Wnt-mediated craniofacial and brain development. Dev. Dyn. 240, 365–371 (2011).

24. C Bänziger, et al., Wntless, a Conserved Membrane Protein Dedicated to the Secretion of Wnt Proteins from Signaling Cells. Cell 125, 509–522 (2006).

25. K Bartscherer, N Pelte, D Ingelfinger, M Boutros, Secretion of Wnt Ligands Requires Evi, a Conserved Transmembrane Protein. Cell 125, 523–533 (2006).

26. R Nygaard, et al., Structural Basis of WLS/Evi-Mediated Wnt Transport and Secretion. Cell (2020).

27. S Gong, et al., A gene expression atlas of the central nervous system based on bacterial artificial chromosomes. Nature 425, 917–925 (2003).

28. YJ Nam, et al., Induction of diverse cardiac cell types by reprogramming fibroblasts with cardiac transcription factors. Dev. (Cambridge) 141, 4267–4278 (2014).

29. LT Shenje, et al., Precardiac deletion of numb and numblike reveals renewal of cardiac progenitors. eLife 2014, e02164 (2014).

30. Q Ma, B Zhou, WT Pu, Reassessment of Isl1 and Nkx2-5 cardiac fate maps using a Gata4-based reporter of Cre activity. Dev. Biol. 323, 98–104 (2008).

31. Y Sun, et al., Islet 1 is expressed in distinct cardiovascula lineages, including pacemaker and coronary vascular cells. Dev. Biol. 304, 286–296 (2007).

32. A Butler, P Hoffman, P Smibert, E Papalexi, R Satija, Integrating single-cell transcriptomic data across different conditions, technologies, and species. Nat. Biotechnol. 36, 411–420 (2018).

33. CS McGinnis, et al., MULTI-seq: sample multiplexing for single-cell RNA sequencing using lipid-tagged indices. Nat. Methods 16, 619–626 (2019).

34. C Hafemeister, R Satija, Normalization and variance stabilization of single-cell RNA-seq data using regularized negative binomial regression. Genome Biol. 20, 296 (2019).

35. T Stuart, et al., Comprehensive Integration of Single-Cell Data. Cell 177, 1888–1902.e21 (2019).

36. T Huynh, L Chen, P Terrell, A Baldini, A fate map of Tbx1 expressing cells reveals heterogeneity in the second cardiac field. Genesis 45, 470–475 (2007).

37. H Xu, et al., Tbx1 has a dual role in the morphogenesis of the cardiac outflow tract. Development 131, 3217–3227 (2004).

38. X Qiu, et al., Single-cell mRNA quantification and differential analysis with Census. Nat. Methods 14, 309–315 (2017).

39. K Van den Berge, et al., Trajectory-based differential expression analysis for single-cell sequencing data. Nat. Commun. (2020).

40. GL Engelmann, KD Boehm, MC Birchenall-Roberts, FW Ruscetti, Transforming growth factor-beta1 in heart development. Mech. Dev. 38, 85–97 (1992).

41. HL Tan, et al., Nonsynonymous variants in the SMAD6 gene predispose to congenital cardiovascular malformation. Hum. Mutat. 33, 720–727 (2012).

42. G Rossi, et al., Capturing Cardiogenesis in Gastruloids. Cell Stem Cell (2020).

## References

1. Y Saga, et al., MesP1 is expressed in the heart precursor cells and required for the formation of a single heart tube. Development 126, 3437–3447 (1999).

2. T Huynh, L Chen, P Terrell, A Baldini, A fate map of Tbx1 expressing cells reveals heterogeneity in the second cardiac field. Genesis 45, 470–475 (2007).

3. S Srinivas, et al., Cre reporter strains produced by targeted insertion of EYFP and ECFP into the ROSA26 locus. BMC Dev. Biol. 1, 1–8 (2001).

4. EG Stanley, et al., Efficient cre-mediated deletion in cardiac progenitor cells conferred by a 3’UTR-ires-Cre allele of the homeobox gene Nkx2-5. Int. J. Dev. Biol. 46, 431–439 (2002).

5. S Gong, et al., A gene expression atlas of the central nervous system based on bacterial artificial chromosomes. Nature 425, 917–925 (2003).

6. YJ Nam, et al., Induction of diverse cardiac cell types by reprogramming fibroblasts with cardiac transcription factors. Dev. (Cambridge) 141, 4267–4278 (2014).

7. J Fu, HM Ivy Yu, T Maruyama, AJ Mirando, W Hsu, Gpr177/mouse Wntless is essential for Wnt-mediated craniofacial and brain development. Dev. Dyn. 240, 365–371 (2011).

8. L Madisen, et al., A robust and high-throughput Cre reporting and characterization system for the whole mouse brain. Nat. Neurosci. 13, 133–140 (2010).

9. LT Shenje, et al., Precardiac deletion of numb and numblike reveals renewal of cardiac progenitors. eLife 2014, e02164 (2014).

10. CS McGinnis, et al., MULTI-seq: sample multiplexing for single-cell RNA sequencing using lipid-tagged indices. Nat. Methods 16, 619–626 (2019).

11. B Bushnell, J Rood, E Singer, BBMerge – Accurate paired shotgun read merging via overlap. PLoS ONE 12, 1–15 (2017).

12. P Melsted, et al., Modular and efficient pre-processing of single-cell RNA-seq. bioRxiv, 1–18 (2019).

13. C Hafemeister, R Satija, Normalization and variance stabilization of single-cell RNA-seq data using regularized negative binomial regression. Genome Biol. 20, 296 (2019).

14. M Tang, et al., Evaluating single-cell cluster stability using the Jaccard similarity index. bioRxiv, 2020.05.26.116640 (2020).

15. BS Mankoo, et al., The concerted action of Meox homeobox genes in required upstream of genetic pathways essential for the formation, patterning and differentiation of somites. Development 130, 4655–4664 (2003).

16. JA Blake, MR Ziman, Pax genes: Regulators of lineage specification and progenitor cell maintenance. Dev. (Cambridge) 141, 737–751 (2014).

17. B Christ, R Huang, J Wilting, The development of the avian vertebral column (2000).

18. U Deutsch, GR Dressler, P Gruss,Pax 1, a member of a paired box homologous murine gene family, is expressed in segmented structures during development. Cell 53, 617–625 (1988).

19. C Ebensperger, et al., Pax-1, a regulator of sclerotome development is induced by notochord and floor plate signals in avian embryos. Anat. Embryol. 191, 297–310 (1995).

20. LM Galli, et al., Identification and characterization of subpopulations of Pax3 and Pax7 expressing cells in developing chick somites and limb buds. Dev. Dyn. 237, 1862–1874 (2008).

21. MD Goulding, G Chalepakis, U Deutsch, JR Erselius, P Gruss, Pax-3, a novel murine DNA binding protein expressed during early neurogenesis. The EMBO J. 10, 1135–1147 (1991).

22. TY de Soysa, et al., Single-cell analysis of cardiogenesis reveals basis for organ-level developmental defects. Nature 572, 120–124 (2019).

23. VM Christoffels, et al., Tbx18 and the fate of epicardial progenitors. Nature 458, E8–E9 (2009).

24. A von Gise, et al., WT1 regulates epicardial epithelial to mesenchymal transition through *β*-catenin and retinoic acid signaling pathways. Dev. Biol. 356, 421–431 (2011).

25. S Conway, J Molkentin, Periostin as a Heterofunctional Regulator of Cardiac Development and Disease. Curr. Genomics 9, 548–555 (2008).

26. P Camelliti, TK Borg, P Kohl, Structural and functional characterisation of cardiac fibroblasts (2005).

27. P Agarwal, et al., Tbx5 is essential for forelimb bud initiation following patterning of the limb field in the mouse embryo (2003).

28. C Rallis, et al., Tbx5 is required for forelimb bud formation and continued outgrowth (2003).

29. L Taher, et al., Global Gene Expression Analysis of Murine Limb Development. PLoS ONE 6, e28358 (2011).

30. SF Gilbert, Formation of the Limb Bud in Developmental Biology. (Sinauer Associates), 6th editio edition, (2000).

31. A Marcil, É Dumontier, M Chamberland, SA Camper, J Drouin, Pitx1 and Pitx2 are required for development of hindlimb buds (2003).

32. E Hornstein, et al., The microRNA miR-196 acts upstream of Hoxb8 and Shh in limb development. Nature 438, 671–674 (2005).

33. CE Nelson, et al., Analysis of Hox gene expression in the chick limb bud, Technical report (1996).

34. A Lother, et al., Cardiac endothelial cell transcriptome. Arter. Thromb. Vasc. Biol. 38, 566–574 (2018).

35. M Ashburner, et al., Gene ontology: Tool for the unification of biology (2000).

36. TheGeneOntologyConsortium, The Gene Ontology Resource: 20 years and still GOing strong. Nucleic acids research 47 (2019).

37. F Supek, M Bošnjak, N Škunca, T Šmuc, REVIGO Summarizes and Visualizes Long Lists of Gene Ontology Terms. PLoS ONE 6, e21800 (2011).

